# Honey bees (*Apis mellifera*) decrease the fitness of plants they pollinate

**DOI:** 10.1101/2023.04.26.538464

**Authors:** Dillon J. Travis, Joshua R. Kohn

## Abstract

Most flowering plants require animal pollination and are visited by multiple pollinator species. Historically, the effects of pollinators on plant fitness have been compared using the number of pollen grains they deposit, and the number of seeds or fruits produced following a visit to a virgin flower. While useful, these methods fail to consider differences in pollen quality and the fitness of zygotes resulting from pollination by different floral visitors. Here we show that, for three common native self-compatible plants in Southern California, super-abundant, non-native honey bees (*Apis mellifera* L.) visit more flowers on an individual before moving to the next plant compared to the suite of native insect visitors. This likely increases the transfer of self-pollen. Offspring produced after honey bee pollination have similar fitness to those resulting from hand self-pollination and both are far less fit than those produced after pollination by native insects or by cross-pollination. Because honey bees often forage methodically, visiting many flowers on each plant, low offspring fitness may commonly result from honey bee pollination of self-compatible plants. To our knowledge, this is the first study to directly compare the fitness of offspring resulting from honey bee pollination to that of other floral visitors.

## Introduction

Approximately 85% of all angiosperms require animal visitation to successfully reproduce [1,2] and flowers of the great majority of these plants are visited by multiple pollinator species [3–5]. It has long been recognized that pollinators can vary in the amount of pollen deposited, the number of seeds produced, or the probability of fruit set resulting from of their visits [6–9]. Far less attention has been paid to differences among pollinators in the fitness of the zygotes that result from the pollen they deposit. This is despite the fact that pollinators vary considerably in a behavior that may strongly affect fitness: the number of flowers on a plant they visit before moving to another plant. Successive visits to flowers on the same plant (geitonogamous visitation) is often the primary cause of self-pollination in plants with large floral displays [10–12]. For self-compatible plants, self-fertilization often severely reduces the fitness of offspring compared to those produced via cross-fertilization [13,14]. For self-incompatible plants, self-pollination may result in fewer seeds set or fruits produced even when stigmatic pollen loads are abundant [15,16].

Inbreeding depression (IBD), the reduced fitness of self-fertilized progeny in comparison to cross-fertilized ones, has been widely measured in plants since it is thought to be a primary selective force acting on plant mating systems [13,17]. Studies of IBD cataloged in several reviews have shown that the fitness of self-fertilized zygotes is often less than half that of cross-fertilized ones [14,18,19]. In addition, reductions in fitness due to self-fertilization tend to be larger for longer-lived plants, reflecting either higher levels of deleterious somatic mutations [20], higher population levels of outcrossing which maintains more genetic load in populations, or larger cumulative effects of deleterious alleles over longer lifespans [21]. Finally, the timing of the expression of IBD varies, with longer-lived, generally more outcrossing plants showing more IBD early in the life cycle (i.e., between fertilization and seed set or germination) than is observed in annual herbs [18, 22].

We were motivated to investigate the fitness of seeds produced by native plants resulting from different pollinators in San Diego County, California, USA for the following reasons. First, studies in this area have shown that non-native, primarily feral, honey bees (*Apis mellifera* L.) are by far the most common floral visitors to native plants in the region [23]. In intact habitats with native vegetation, honey bees make up approximately 75% of all floral visitors and are even more dominant on the most abundantly blooming species, often exceeding 90% of all visitors [23]. This is among the highest levels of honey bee community dominance recorded anywhere in the world [24]. At the same time, San Diego County is a biodiversity hotspot with over 600 species of native bees [25] and at least 2,400 plant taxa [26], so the impacts of this highly abundant non-native pollinator are of interest. Second, honey bees forage methodically, and casual observation suggested that their levels of geitonogamous visitation are perhaps higher than the average among native pollinating insects. Self-fertilization rates are known to increase as the number of flowers consecutively visited on a plant during a foraging bout increase [27]. This led us to hypothesize that if honey bees make more geitonogamous visits, the offspring that result from their pollination might have reduced fitness.

## Results and Discussion

We studied three common and abundantly blooming native plants: *Salvia mellifera* Greene and *S. apiana* Jeps., which are perennial shrubs and *Phacelia distans* Benth., an annual herb. All three species are self-compatible and produce large inflorescences with multiple flowers open simultaneously. Honey bees on average made 1.76 - 2.33 times as many geitonogamous visits per plant compared to non-*Apis* floral visitors which comprise a suite of overwhelmingly native insects [25] (p < 0.0001 for all three species, Figure 1, Table S4).

**Figure 1.**
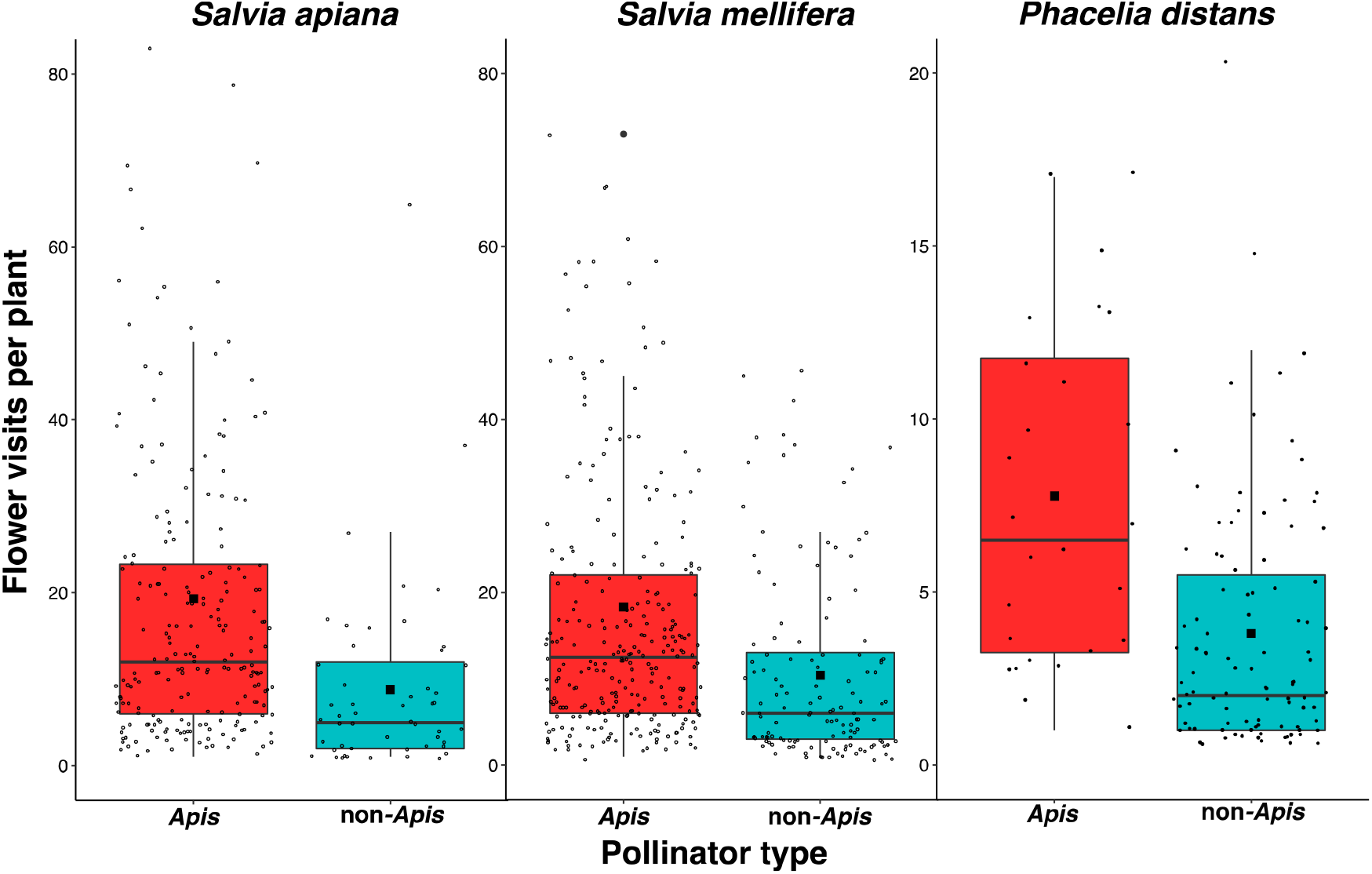
The number flowers visited per plant by *Apis* and non-*Apis* pollinators during single foraging bouts. Boxplots: the lines show medians; black boxes show means; boxes and the whiskers represent interquartile ranges; dots represent individual insect foraging bouts. For all three species, honey bees visited more flowers per plant compared to non-*Apis* insects (P < 0.0001). See Table S4 for statistical model outputs.

All three plant species exhibited strong inbreeding depression, with cross-pollination providing 2 to 10-fold higher multiplicative fitness values when germinated and then grown in a greenhouse (Figure 2; Tables S1-3, S6). Consistent with results of meta-analyses of inbreeding depression [28,29], the two perennial species (*S. mellifera* and *S. apiana*) exhibited greater fitness differences between self- and cross-pollination treatments than the annual *P. distans*. We also observed strong inbreeding depression in both early (seed set, germination, Tables S7 & S8) and later (number of leaves, Table S10) life stages in the *Salvia* species, while differences between self- and cross-pollination treatments in the annual *P. distans* were significant but smaller than the perennial species for seed set and germination rate, and there were differences among these treatments in the production of flowers (Tables S3 & S11). For all three species, there was no significant difference among treatments in seedling survival to ten weeks (Table S9).

**Figure 2.**
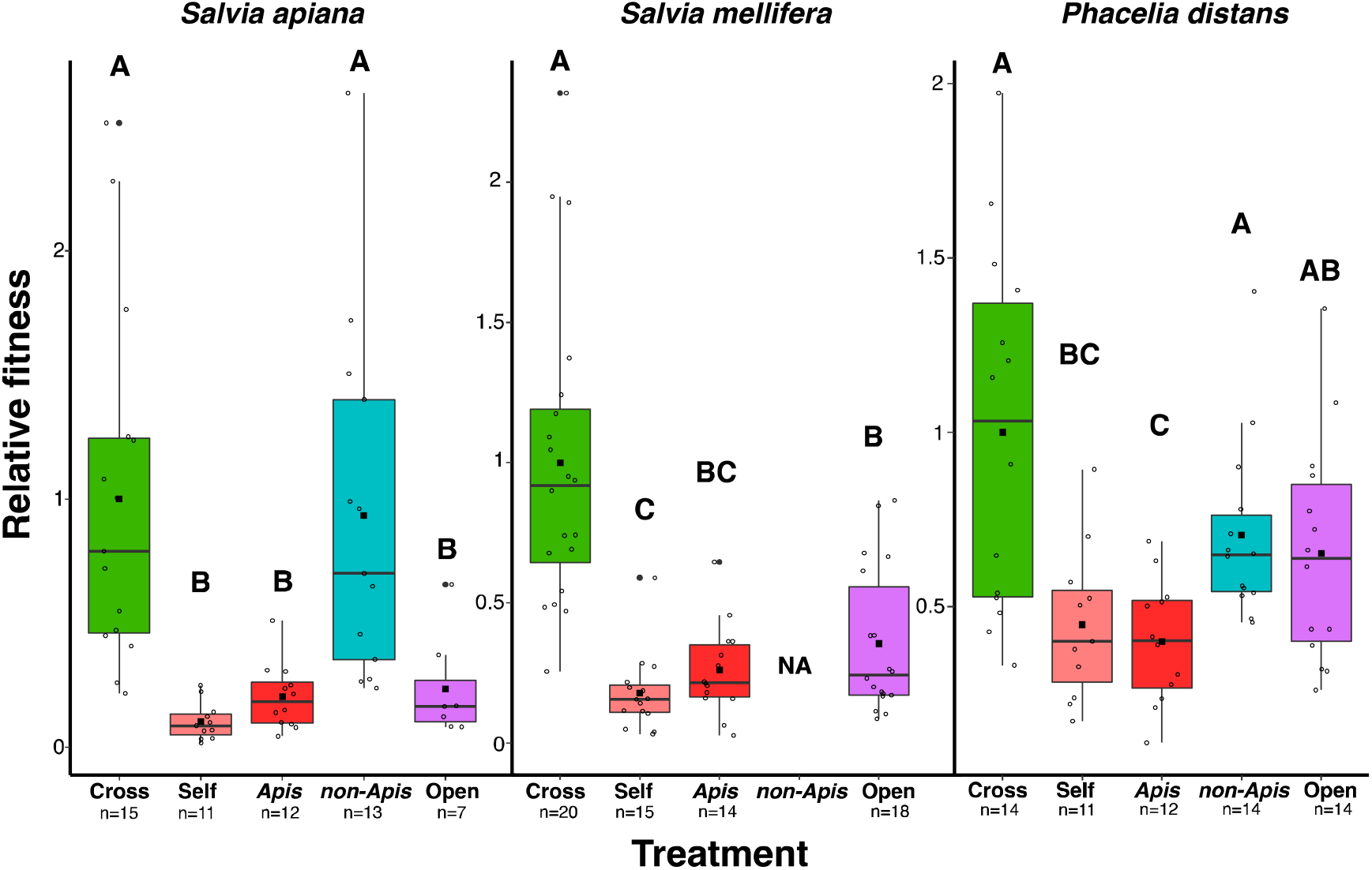
Relative fitness of each pollination treatment for each plant species. All values have been standardized so that the mean fitness of cross treatments equal 1. Boxplots: the lines show medians; black boxes show means; boxes and the whiskers represent interquartile ranges; dots represent mean values for maternal plant families. Bold letters signify statistical differences between treatments. Pollination treatment had a significant effect for all three species (P < 0.0001). See Tables S1-3 for full model outputs for each fitness trait measured.

Fitness resulting from single *Apis* visits did not differ from the hand self-pollination treatment for all three species. For the two species (*S. apiana* and *P. distans*) with sufficient native insect visitation to allow us to assess the reproductive consequences, fitness resulting from single visits by native pollinators was not different than fitness from hand cross-pollination and two to tenfold greater than fitness resulting from pollination by honey bees or hand self-pollination. For the two species of *Salvia*, the fitness measured for open-pollinated seeds was not significantly different from the fitness of seeds from the *Apis* pollination treatment, likely reflecting the fact that >90% of visits to those species were by honey bees (Table S12). For *P. distans*, open-pollinated seeds had an average fitness intermediate between self- and cross-treatments and intermediate between non-*Apis* and *Apis* pollination treatments. This may reflect the sizeable fraction (20-25%) of visitors to this species that were native pollinators, or that *P. distans* plants have significantly fewer flowers open simultaneously compared to the *Salvia* species. It should be noted that these fitness measures come from greenhouse estimates of fitness components. Greenhouse experiments often underestimate the cost of selfing relative to outcrossing in comparison to estimates based on measurements made in more stressful, field environments [30,31].

Differences between *Apis* and non-*Apis* pollination treatments in seed set could reflect pollen limitation if single honey bee visits deliver fewer pollen grains than those of native insects, and if the amount of pollen they deliver limits seed set. We measured stigmatic pollen loads following single visits to previously unvisited flowers by honey bees and native pollinators. For *S. apiana* and *P. distans*, pollen deposition following single visits by honey bees and native insects did not differ significantly, though honey bees deposited somewhat fewer grains, on average, in both species (Figure 3, Table S5). Whether the amount of pollen deposited by single honey bee visits limited seed set is more difficult to assess. All three species have four ovules per flower and can produce a maximum of 4 seeds per fruit. When given abundant cross-pollen, all three species averaged approximately 2 seeds per flower (Tables S1-3). The mean single visit pollen deposition by honey bees was 7.9, 19.8, and 29.9 grains for *S. apiana, S. mellifera* and *P. distans* respectively, which seems to be sufficient for full seed set since ad-lib deposition of self-pollen did not produce significantly more seeds than single *Apis* visits in any of the three species. For *Salvia apiana* and *S. mellifera*, the open pollination treatment, in which flowers are typically visited multiple times per day by honey bees (D.T. unpublished data) also did not produce significantly more seeds than did single honey bee visits. For *P. distans*, which garners more visitation from native insects, open pollinated seed set did not differ from that which resulted from hand cross-pollination (Table S3). In contrast, for both *S. apiana* and *P. distans*, single visits by non-*Apis* pollinators resulted in seed set values that did not differ from application of abundant cross-pollen, indicating differences in the quality of pollen delivered between the two types of pollinators. Further, in *S. apiana*, the species with the lowest pollen deposition following single honey bee visits, differences between *Apis* and non-*Apis* pollination treatments for fitness components that occur after seed set (germination and leaf production) are significant, again indicating differences in the quality of pollen received from different pollinators (Table S1).

**Figure 3.**
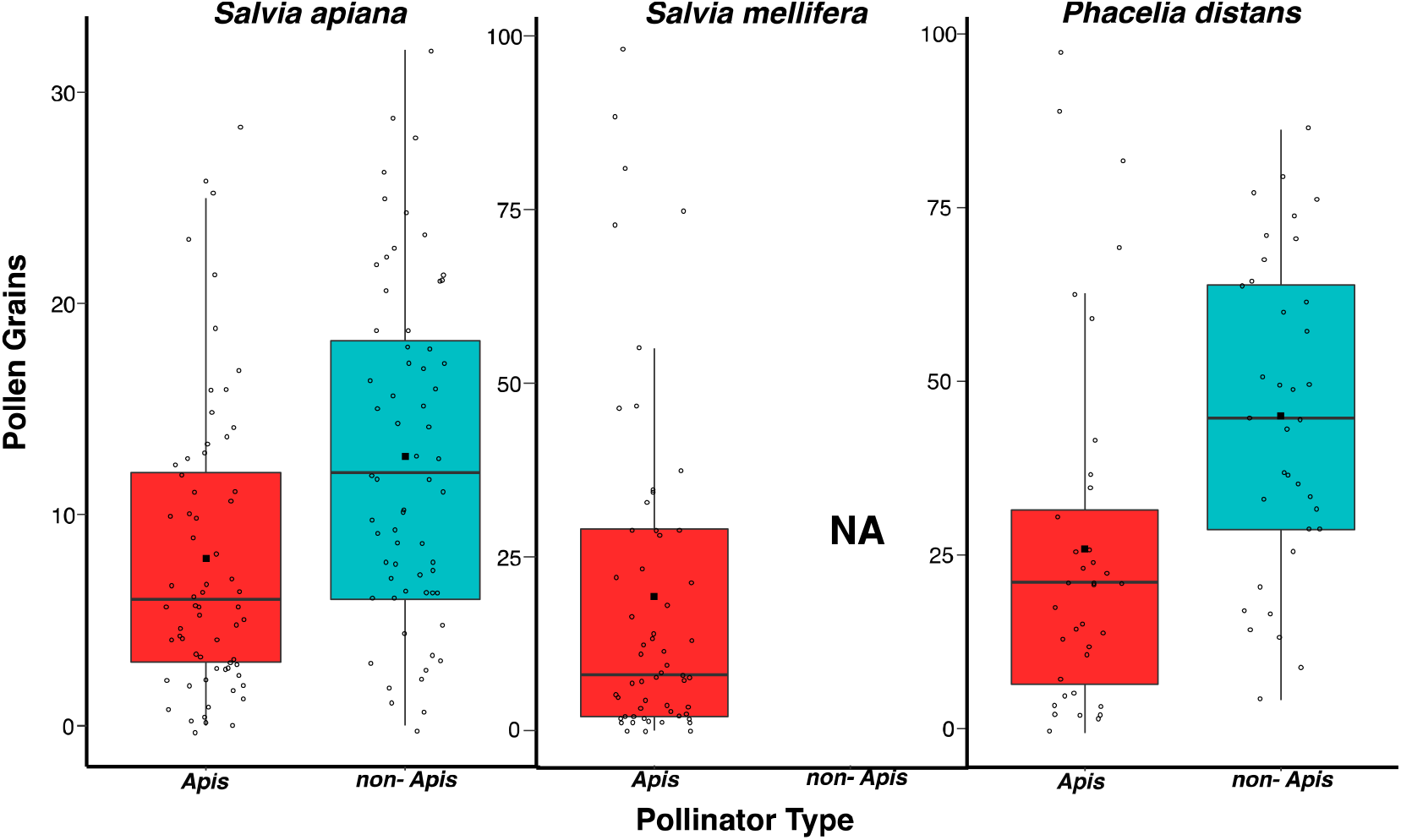
The number of pollen grains deposited on stigmas in a single visit by *Apis* and non-*Apis* pollinators. Boxplots: features are the same as in Figure 1. Honey bees deposit a similar number of pollen grains as non-*Apis* insects for *S. apiana* and *P. distans* (P > 0.05). See Table S5 for full model outputs.

Our findings have several major implications. First, honey bees are estimated to be the most frequent floral visitors to natural vegetation worldwide, accounting for 13% of all floral visits globally [24]. Pollination studies of many plant species, or whole-community pollination networks, often use visitation rates as the measure of a pollinator’s importance. Some go further and refine these estimates of importance by multiplying them by single visit pollinator effectiveness, measured as pollen grains delivered, seeds set, or probability of fruit set [16,32,33]. While measures of single-visit effectiveness of honey bees vary across plant species, meta-analyses have shown that, on average, their effectiveness is not different than the mean of other pollinators visiting the same species [24,34]. But if pollinators deliver pollen of different quality that leads to strong differences in the fitness of the offspring produced, this will impact the importance of different pollinators to a plant’s reproductive success. The fitness differences resulting from honey bee versus primarily native, non-*Apis* insect visits measured here indicate that a pollinator’s real importance may be strongly influenced by their foraging behavior (e.g. differences in geitonogamous visitation) and the pollen quality delivered. High levels of self-pollen delivered by honey bees may also help explain why, across a wide range of 41 crops, increased visitation by non-honey bee insects increased seed or fruit set regardless of the amount of honey bee visitation [35].

Few studies have measured the fitness of seeds resulting from different pollinators or sets of pollinators [8, 36–38], and none have directly compared the fitness effects of honey bee visits to those of other insects. In the most relevant study, Herrera [8] planted seeds of *Lavandula latifolia* that resulted from exposure to pollinators at different times of day. In a field planting study, seeds from flowers exposed only to pollinators during the early morning and late evening were significantly less likely to germinate and survive than seeds which resulted from flowers exposed to pollinators only during the middle of the day. This plant has an array of floral visitors. Large bees, which were predominantly honey bees, visit throughout the day, but lepidoptera and small bees, which made up a minority of pollinators, were more common in the middle of the day than when cooler temperatures prevailed. Herrera [8] attributed the observed fitness differences to the fact that small bees and, particularly, lepidoptera, make fewer geitonogamous visits than do large bees. The fact that both Herrera’s study and ours implicate honey bees as pollinators whose services result in low quality offspring should motivate further research into the generality of whether honey bees tend to deliver more geitonogamous self-pollen than other pollinators, and the fitness consequences that may result.

In San Diego County (USA), non-native, feral, honey bees dominate the pollinator community that visits native plants. Many of these native plant species are self-compatible, so if honey bees generally transfer pollen that reduces the fitness of seeds produced there could be many ecological consequences. We speculate on just three. First, if honey bees generally lower seed fitness of native plants, this could make the native the plant community more susceptible to invasion by introduced plant species that do not require insect pollination, or which are historically highly self-fertilizing, such as invasive annual Mediterranean grasses and mustard species (*Hirschfeldia incana* and *Brassica nigra*) that currently occupy much of the space between native shrubs and increase the ecosystem’s susceptibility to fire. Second, to the degree that individual species become more inbred due to honey bee pollination, their evolutionary future might be compromised. This is important because San Diego County is biodiversity hotspot having the most plant taxa of any county in the USA [26]. Third, to the extent that honey bees focus their resource gathering efforts on the most abundantly blooming taxa, ignoring, or at least not fully dominating visitation to rarer taxa [23], they might preserve plant diversity by reducing the fitness of the most abundant plant species. It is impossible to predict all the repercussions of high honey bee abundance and the lower reproductive fitness of the plants they pollinate, but these effects are likely to be substantial.

## Materials and Methods

We compared the effects of honey bees and native pollinating insects on the reproductive fitness of three common plant species in coastal sage scrub habitats of San Diego County: *Phacelia distans* (common phacelia, Boraginaceae) an annual herb, *Salvia mellifera* (black sage, Lamiaceae) and *Salvia apiana* (white sage, Lamiaceae) which are perennial shrubs. All three species produce hermaphroditic flowers on multiple inflorescences and commonly display dozens to hundreds of flowers at once, with both pollen producing and pollen receptive flowers open simultaneously. All three species are self-compatible [39,40] and therefore visits to multiple flowers on the same plant may lead to geitonogamous self-fertilization and potentially lower the fitness of the resulting zygotes. The two *Salvia* species are large shrubs, and during peak bloom, are often the most abundantly flowering species at a given site and attract a large (>90%) fraction of floral visits from honey bees. *Phacelia distans*, an annual, is rarely the most abundantly blooming plant at a given site and a greater proportion of flower visitors are non-*Apis* insects compared to the two *Salvia* species (Table S12).

To determine if non-native honey bees make more geitonogamous visits than native pollinators, in 2018 we identified sites where at least one of our plant species occurred in intact coastal sage scrub habitat (*S. apiana*-4 sites, *S. mellifera*-2 sites, *P. distans*-1 site). During peak bloom for each species, we collected visitation data from 10:00 – 16:00, on days with less than 50% cloud cover, little to no wind, and air temperatures exceeding 16°C to minimize environmental impacts on foraging behavior. Visitation data were collected by observing a pollinator approach a plant and counting the number of flowers they visited before moving on to another plant or out of our field of vision. For each foraging bout, we recorded the site, date, plant species and individual, pollinator type (*Apis* or non-*Apis*), and the number of flowers the pollinator visited before moving on. Due to the diversity of native insect visitors, our interest in comparing non-native honey bees to the native pollinator community, and the expertise required to identify insects on the fly to the species level, pollinators were classified as *Apis* or non*-Apis*. We recorded the number of flower visits per plant for 212 honey bees and 51 non-*Apis* visitors on *S. apiana*, 274 honey bees and 131 non-*Apis* visitors on *S. mellifera*, and 27 honey bees and 99 non-*Apis* visitors on *P. distans*. Detailed lists and frequencies of insect visitors to plants in western San Diego County can be found at https://library.ucsd.edu/dc/collection/bb0072854b.

We measured the fitness effects of different pollination treatments as follows. In the spring of 2018, we identified 5 individuals of *S. apiana* at each of 3 sites, and in the spring of 2019, we added 15 individuals of *S. apiana* at each of 2 sites, 15 *S. mellifera* individuals at each of 2 sites and 30 *P. distans* individuals at a single site (Table S13). Before pollination treatments, we placed mesh pollinator-exclusion bags over 6 inflorescences on each plant to prevent visitation. During peak bloom for each species, we returned to individual plants and removed the mesh bags to expose unvisited female phase flowers to one of six pollination treatments: 1. Open-pollinated (control) flowers were exposed to visitation for the duration of a flower’s life to assess pollination and seed set in field conditions. 2. Cross-pollinated flowers were hand pollinated using pollen from 3-5 individuals at least 20 meters away to minimize any impact of donor identity. Pollen was transferred to a stigma using forceps. 3. Self-pollinated flowers were hand pollinated with pollen acquired from 3-5 fresh anthers from the same plant. 4. Honey bee-pollinated flowers were exposed until a single honey bee was observed foraging on an unvisited flower, after which the inflorescence was bagged to exclude further visitation. 5. Non-*Apis* insect-pollinated flowers were exposed until a single non-*Apis* insect visited a flower and then bagged to exclude further visitation. 6. The pollinator exclusion treatment in which flowers where enclosed in mesh bags for their lifetime to determine the degree to which pollinators are necessary for set seed. The calyx of each treatment flower was marked with a dot of acrylic paint, and after 4 to 6 weeks mature seeds were recovered. Due to insufficient visitation by non-*Apis* insects to experimental flowers of *Salvia mellifera*, we were unable to assess the impact non-*Apis* pollination for this species.

To assess single visit pollen deposition, we exposed receptive, unvisited stigmas to one visit from a honey bee or a non-*Apis* insect. After the insect contacted the stigma, we immediately collected stigmas and pressed them into fuchsin jelly on a microscope slide and later counted pollen grains with a dissecting microscope. We collected 64 *Apis* and 63 non-*Apis* pollinated stigmas from *S. apiana*, and 37 *Apis* and 35 non-*Apis* pollinated stigmas from *P. distans*. For *S. mellifera*, we collected 55 stigmas that were visited by honey bees.

We counted the number of seeds produced for each treatment for every maternal plant. Seeds that were <10% normal size and appeared aborted were not included. After counting, seeds were washed with a 10% bleach solution, dried, and stored at 4°C for at least 6 months to help break seed dormancy. In late January 2020, seeds were removed from storage and placed into sterile filter paper lined petri dishes. Seeds produced from the pollinator exclusion treatment were not included due to greenhouse space constraints. Since both *Salvia* species have low (10-20%) germination rates [38], we employed a variety of treatments to promote germination. After being removed from 4°C, *Salvia* seeds were heated to 70°C for 1 hour [41], treated with a 1:10 dilution of Wrights™ liquid smoke [42], then soaked in 5-6 ml of a 500ppm solution of gibberellic acid in deionized water overnight [43]. To further stimulate germination, petri dishes containing the hydrated *Salvia spp*. seeds were placed on a warming mat set to 26°C and exposed to the natural light in the greenhouse [41]. *P. distans* seeds germinate readily so the seeds were placed in petri dishes, soaked in 5-6 ml of deionized water, and placed in the dark to encourage germination.

Each morning we examined all petri dishes, if a seed’s radical had emerged, the date was recorded, and the seed was planted in a pot (5 × 12” tree pots, Stuewe & Sons Inc.) filled with native topsoil and placed in a randomized position within our greenhouse. Due to the space limitations, if two seeds from the same parent and treatment germinated on the same day, they were placed on opposite sides of the same pot. After 4 weeks, if both seedlings were alive in the same pot, one was culled at random. After 4 weeks we counted the number of seeds that failed to germinate in each petri dish.

Seedlings were watered with 200 ml of water twice a week using an automated drip irrigation system (DripWorks, Inc.). All pot positions were re-randomized within the greenhouse 4 and 8 weeks after the beginning of the experiment to reduce microclimate effects. 10 weeks after germination the following were measured: percent germination, survival to 10 weeks, and for *Salvia apiana* and *S. mellifera*, the number of leaves at 10 weeks as a proxy for size [18,44] which is strongly associated with reproductive output in plants [45]. For the annual *P. distans*, which reaches sexual maturity several weeks after germination, we counted the number of flowers each plant produced. We then calculated mean values of these traits for each maternal parent plant for every treatment. Relative fitness for each treatment and maternal plant was calculated using a multiplicative fitness function [18,44]. For *S. apiana* and *S. mellifera*, relative fitness of each treatment was calculated as the product of mean seed set, germination success, proportion of seedlings that survived at 10 weeks, and the number of leaves at 10 weeks. For *P. distans*, relative fitness was calculated in a similar fashion except the mean number of flowers produced for each treatment was used in place of the number of leaves.

## Statistical Analysis

All statistical analyses were conducted in R (v. 3.5.0, 2021), using packages lme4 [46], nlme [47], lmerTest [48], lattice [49], Rmisc [50], multcomp [51], lsmeans [52], ggplot2 [53], and plyr [54].

We used linear mixed-effect models (LMM) to determine differences in geitonogamous visitation and single visit pollen deposition between *Apis* and non-*Apis* pollinators for each plant species. Random effects considered were date, individual plant identity, year, and site when applicable. Individual plant identity was nested in site, and site was nested in date. The fixed effect considered in both models was pollinator type (*Apis* or non-*Apis*). The number of visits per plant was log_10_ transformed and single visit pollen deposition was square root transformed to fit model assumptions. To assess differences in relative fitness (log_10_ transformed) between pollination treatments, we constructed a LMM with maternal identity, site, and year as random effects when applicable for each plant species. Again, maternal identity was nested in site, and site was nested in year. The fixed effect considered was pollination treatment in all models. Random effects in each model were tested by performing likelihood ratios tests. If models failed to converge, the random effect that caused the failure was removed.

## Acknowledgments

D. Meszaros, J. Bohey, J. Noonan, N. Callen, B. Tsai, M. Musse, and J. Waits assisted in data collection. D.T. was supported by grants from Sea and Sage Audubon Society, the Messier Family Fund and by a UC MRPI grant to J.R.K. This work was performed in part at the University of California Natural Reserve System’s Elliott Chaparral and Dawson Los Monos Canyon Reserves.

