## Supplemental Tables 1-13 for "Honey bees (*Apis mellifera*) decrease the fitness of plants they pollinate"

**This PDF file includes:**

Tables S1 to S13

**Supplementary Tables**

**Table S1. Summary of fitness trait values for *S. apiana* pollination treatments**. Values in parentheses are 1 standard deviation. Superscript letters indicate significant differences between treatments determined with least squares means tests with sidak adjusted significance levels for 5 or 6 families ($\propto$ = 0.05).

| ***Salvia apiana*** |  |  | **Pollination** | **Treatment** |  |  |
| --- | --- | --- | --- | --- | --- | --- |
| **Trait** | **Cross** | **Self** | ***Apis*** | **non-*Apis*** | **Open** | **Exclusion** |
| **Seed set** | 1.85**^A^** (0.66) | 0.92**^B^**  (0.53) | 0.85**^B^** (0.41) | 1.74**^A^** (0.79) | 0.86**^B^** (0.72) | 0.54**^B^** (0.51) |
| **Germination**  **Success (%)** | 59.29**^A^** (29.1) | 18.11**^BC^**  (21.1) | 29.08**^B^** (29.7) | 51.69**^A^** (31.6) | 15.21**^C^** (16.4) | NA |
| **Survival**  **(%)** | 58.33**^A^** (38.9) | 54.52**^A^**  (39.4) | 45.71**^A^** (20.9) | 69.76**^A^** (32.2) | 50.0**^A^** (47.7) | NA |
| **Number of**  **leaves at 10 wks** | 22.81**^A^** (6.09) | 12.24**^C^**  (5.47) | 12.19**^C^** (3.61) | 19.25**^AB^** (3.89) | 14.14**^BC^** (4.14) | NA |
| **Relative**  **Fitness** | 1.00**^A^** (0.71) | 0.11**^B^**  (0.08) | 0.20**^B^** (0.13) | 0.93**^A^** (0.71) | 0.24**^B^** (0.21) | NA |

**Table S2. Summary of fitness trait values for *S. mellifera* pollination treatments.** Superscript letters indicate significant differences between treatments determined with least squares means tests with sidak adjusted significance levels for 4 or 5 families ($\propto$ = 0.05).

| ***Salvia mellifera*** |  |  | **Pollination** | **Treatment** |  |  |
| --- | --- | --- | --- | --- | --- | --- |
| **Trait** | **Cross** | **Self** | ***Apis*** | **non-*Apis*** | **Open** | **Exclusion** |
| **Seed set** | 2.08**^A^**  (0.49) | 1.04**^BC^**  (0.48) | 0.83**^C^** (0.49) | NA | 1.41**^B^** (0.48) | 0.32**^D^** (0.34) |
| **Germination Success (%)** | 70.03**^A^** (25.4) | 45.86**^B^**  (25.5) | 47.87**^B^** (25.6) | NA | 45.95**^B^** (27.6) | NA |
| **Survival**  **(%)** | 49.27**^A^** (29.9) | 38.45**^A^**  (38.6) | 34.96**^A^** (32.78) | NA | 48.13**^A^** (36.0) | NA |
| **Number of**  **leaves at 10 wks** | 25.38**^A^** (9.01) | 13.07**^B^**  (4.61) | 17.81**^B^** (6.31) | NA | 16.24**^B^** (6.31) | NA |
| **Relative**  **Fitness** | 1.00**^A^**  (0.55) | 0.18**^C^**  (0.14) | 0.26**^BC^** (0.16) | NA | 0.35**^AB^** (0.26) | NA |

**Table S3. Summary of fitness trait values for *P. distans* pollination treatments.** Superscript letters indicate significant differences between treatments determined with least squares means tests with sidak adjusted significance levels for 5 or 6 families ($\propto$ = 0.05).

| ***Phacelia distans*** |  |  | **Pollination** | **Treatment** |  |  |
| --- | --- | --- | --- | --- | --- | --- |
| **Trait** | **Cross** | **Self** | ***Apis*** | **non-*Apis*** | **Open** | **Exclusion** |
| **Seed Set** | 2.12**^A^**  (1.0) | 1.65**^AB^**  (0.7) | 1.38**^BC^** (0.58) | 2.18**^A^**  (0.7) | 2.05**^AB^** (0.7) | 0.92**^C^** (0.7) |
| **Germination Success (%)** | 84.7**^A^** (17.6) | 70.0**^B^**  (25.7) | 80.4**^AB^** (24.2) | 86.4**^AB^**  (12.4) | 82.4**^AB^** (0.15) | NA |
| **Survival**  **(%)** | 100**^A^**  (0) | 96.9**^A^**  (10.1) | 100**^A^**  (0) | 97.4**^A^**  (6.7) | 100**^A^**  (0) | NA |
| **Flower**  **Number** | 391.6**^A^**  (80) | 256.1**^B^**  (60) | 241.7**^B^**  (39) | 248.6**^B^**  (71) | 277.1**^B^** (55) | NA |
| **Relative**  **Fitness** | 1.00**^A^** (0.52) | 0.45**^BC^**  (0.22) | 0.40**^C^** (0.18) | 0.71**^A^** (0.26) | 0.65**^AB^** (0.32) | NA |

**Table S4. Geitonogamous Visitation.** Significance levels of variables used to evaluate the effects of Pollinator Type (*Apis* or non-*Apis*) on the number of flowers visited per plant before moving on. Maternal plant identity was nested in site, then nested in date and were included in models as random effects when applicable. Random effects that caused models to fail to converge were removed.

| Model: log_10_(number of visits per plant) ~ Pollinator Type + (1\|Maternal Plant: Site) | | | | | |
| --- | --- | --- | --- | --- | --- |
| **Plant spp.** | **Variable** | **Test Statistic** | | | **p-value** |
| ***Salvia apiana*** | **Pollinator Type** | **F_(1, 157.35)_ = 33.5** | | | **<0.0001*** |
|  | Maternal Identity: Site | $\chi_{1}^{2}$ = 0.373 | | | 0.542 |
| Model: log_10_(number of visits per plant ~ Pollinator Type + (1\| Maternal Identity: Site: Date) | | | | | |
| ***Salvia mellifera*** | **Pollinator Type** | **F_(1, 386.99)_ = 50.03** | **<0.0001*** | | |
|  | **Maternal Identity: Site: Date** | $\boldsymbol{\chi}_{\boldsymbol{1}}^{\boldsymbol{2}}$ **= 53.97** | **<0.0001*** | | |
| Model: log_10_(number of visits per plant ~ Pollinator Type + (1\| Maternal Identity: Date) | | | | | |
| ***Phacelia distans*** | **Pollinator Type** | **F_(1, 122.98)_ = 27.77** | | **<0.0001*** | |
|  | Maternal Identity: Date | $\chi_{1}^{2}$ = 1.84 | | 0.175 | |

**Table S5.** **Single Visit Pollen Deposition.** Significance levels of variables used to evaluate the effects of Pollinator Type (*Apis* or non-*Apis*) on the number of pollen grains deposited on a stigma in a single visit. No model was used for *Salvia mellifera*, as there was insufficient visitation by non-*Apis* pollinators.

| Model: (number of pollen grains)^1/2^ ~ Pollinator Type | | | |
| --- | --- | --- | --- |
| **Plant spp.** | **Variable** | **Test Statistic** | **p-value** |
| ***Salvia apiana*** | Pollinator Type | F_(1, 1.32)_ = 0.55 | 0.568 |
| ***Salvia mellifera*** | Pollinator Type | NA | NA |
| Model: number of pollen grains ~ Pollinator Type | | | |
| ***Phacelia distans*** | Pollinator Type | F_(1, 1.31)_ = 1.31 | 0.392 |

**Table S6. Relative Fitness**. Significance levels of variables used to evaluate the effects of Pollination Treatment on the relative fitness of seedlings. Maternal plant identity was nested in Site, then nested in Year, and they were included in models as random effects when applicable. If a random effect caused a model to fail to converge, it was removed.

| Model: log_10_(relative fitness) ~ Treatment + (1\|Maternal Identity: Site: Year) | | | |
| --- | --- | --- | --- |
| **Plant spp.** | **Variable** | **Test Statistic** | **p-value** |
| ***Salvia apiana*** | **Treatment** | **F_(4, 43.67)_ = 25.275** | **<0.0001*** |
|  | Maternal Identity: Site:Year | $\chi_{1}^{2}$ = 2.08 | 0.149 |
| Model: log_10_(relative fitness) ~ Treatment + (1\|Maternal Identity: Site) | | | |
| ***Salvia mellifera*** | **Treatment** | **F_(3, 49.36)_ = 22.879** | **<0.0001*** |
|  | Maternal Identity: Site | $\chi_{1}^{2}$ = 0.03 | 0.854 |
| Model: log_10_ (relative fitness) ~ Treatment +  (1\|Maternal Identity) | | | |
| ***Phacelia distans*** | Treatment | 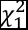**F_(4, 49.4)_ = 8.52** | **<0.0001*** |
|  | Maternal Identity | $\chi_{1}^{2}$ = 3.2 | 0.074 |

**Table S7. Seed Set per flower.** Significance levels of variables used to evaluate the effects of Pollination Treatment on the number of seeds set per flower. Maternal plant identity was nested in Site, then nested in year, and were included in models as random effects when applicable. If a random effect caused a model to fail to converge, it was removed.

| Model: log_10_(seed set) ~ Treatment + (1\|Maternal Identity: Site: Year) | | | |
| --- | --- | --- | --- |
| **Plant Species** | **Variable** | **Test Statistic** | **p-value** |
| ***Salvia apiana*** | **Treatment** | **F_(5, 91.4)_ = 22.09** | **<0.0001*** |
|  | **Maternal Identity: Site: Year** | $\boldsymbol{\chi}_{\boldsymbol{1}}^{\boldsymbol{2}}$ **= 3.89** | **0.0487*** |
| Model: log_10_(seed set) ~ Treatment + (1\|Maternal Identity) + (1\|Site) | | | |
| ***Salvia mellifera*** | Treatment | **F_(4, 137)_ = 54.36** | **<0.0001*** |
|  | Maternal Identity: Site | $\chi_{1}^{2}$ = 0.213 | 0.375 |
| Model: log_10_(seed set) ~ Treatment + (1\|Maternal Identity) | | | |
| ***Phacelia distans*** | **Treatment** | **F_(5, 115)_ = 13.14** | **<0.0001*** |
|  | **Maternal Identity** | $\boldsymbol{\chi}_{\boldsymbol{1}}^{\boldsymbol{2}}$ **= 6.46** | **0.01*** |

**Table S8**. **Germination Probability.** Significance levels of variables used to evaluate the effects of Pollination Treatment on the probability of germination. Maternal plant identity was nested in Site, then nested in year, and were included in models as random effects when applicable. If a random effect caused a model to fail to converge, it was removed.

| Model: Germination ~ Treatment + (1\|Maternal Identity: Site: Year),  binomial, link = logit, optimizer = bobyqa | | | |
| --- | --- | --- | --- |
| **Plant spp.** | **Variable** | **Test Statistic** | **p-value** |
| ***Salvia apiana*** | **Treatment** | $\boldsymbol{\chi}_{\boldsymbol{4}}^{\boldsymbol{2}}$ **= 87.54** | **<0.0001*** |
|  | **Maternal Identity: Site: Year** | $\boldsymbol{\chi}_{\boldsymbol{1}}^{\boldsymbol{2}}$ **= 42.55** | **<0.0001*** |
| Model: Germination ~ Treatment + (1\|Maternal Identity: Site),  binomial, link = logit, optimizer = bobyqa | | | |
| ***Salvia mellifera*** | **Treatment** | $\boldsymbol{\chi}_{\boldsymbol{3}}^{\boldsymbol{2}}$ **= 50.94** | **<0.0001*** |
|  | **Maternal Identity: Site** | $\boldsymbol{\chi}_{\boldsymbol{1}}^{\boldsymbol{2}}$ **= 49.4** | **<0.0001*** |
| Model: Germination ~ Treatment + (1\|Maternal Identity),  binomial, link = logit, optimizer = bobyqa | | | |
| ***Phacelia distans*** | **Treatment** | $\boldsymbol{\chi}_{\boldsymbol{4}}^{\boldsymbol{2}}$ **= 12.96** | **0.011*** |
|  | Maternal Identity | $\chi_{1}^{2}$ = 1.13 | 0.79 |

**Table S9. Survival to 10 weeks.** Significance levels of variables used to evaluate the effects of Pollination Treatment on seedling survival to 10 weeks. Maternal plant identity, site, and year were included in models as random variables when applicable. Maternal Identity was nest in site, then nested in year. If a random effect caused a model to fail to converge, it was removed.

| Model: Survival ~ Treatment + (1\|Maternal Identity: Site: Year), binomial,  link = logit, optimizer = bobyqa | | | |
| --- | --- | --- | --- |
| **Plant spp.** | **Variable** | **Test Statistic** | **p-value** |
| ***Salvia apiana*** | Treatment | $\chi_{4}^{2}$ = 5.62 | 0.23 |
|  | Maternal Identity: Site: Year | $\chi_{1}^{2}$ = 0.58 | 0.75 |
| Model: Survival ~ Treatment + (1\|Maternal Identity), binomial,  link = logit, optimizer = bobyqa | | | |
| ***Salvia mellifera*** | Treatment | $\chi_{3}^{2}$ = 3.92 | 0.27 |
|  | Maternal Identity | $\chi_{1}^{2}$ = 0.03 | 0.90 |
| Model: Survival ~ Treatment + (1\|Maternal Identity), binomial,  link = logit, optimizer = bobyqa | | | |
| ***Phacelia distans*** | Treatment | $\chi_{4}^{2}$ = 4.19 | 0.38 |
|  | Maternal Identity | $\chi_{1}^{2}$ = 5.45 | 0.24 |

**Table S10. Number of leaves at 10 weeks.** Significance levels of variables used to evaluate the effects of Pollination Treatment on the number of leaves present at 10 weeks. Maternal plant identity was nested in Site, then nested in year and were included in models as random variables when applicable. Random effects that caused models to fail to converge were removed.

| Model: # of leaves ~ Treatment + (1\|Year) | | | |
| --- | --- | --- | --- |
| **Plant spp.** | **Variable** | **Test Statistic** | **p-value** |
| ***Salvia apiana*** | **Treatment** | **F_(4, 46)_ = 12.41** | **<0.0001*** |
|  | Year | $\chi_{1}^{2}$ = 0.08 | 0.776 |
| Model: # of leaves ~ Treatment + (1\|Maternal Identity: Site) | | | |
| ***Salvia mellifera*** | **Treatment** | **F_(3, 51.8)_ = 11.63** | **<0.0001*** |
|  | Maternal Identity: Site | $\chi_{1}^{2}$ = 0.59 | 0.439 |

**Table S11. Number of flowers produced.** Significance levels of variables used to evaluate the effects of Pollination Treatment on the number of flowers produced in a plant’s lifetime. Maternal plant identity was included in the model as a random variable.

| Model: Number of Flowers ~ Treatment + (1\|Maternal Identity) | | | |
| --- | --- | --- | --- |
| **Plant spp.** | **Variable** | **Test Statistic** | **p-value** |
| ***Phacelia distans*** | **Treatment** | **F_(4, 59)_ = 11.52** | **<0.0001*** |
|  | Maternal Identity | $\chi_{1}^{2}$ = 0.03 | 0.847 |

**Table S12. *Apis* and non-*Apis* floral visitors for plant species at each site.** Data was collected in 2021 by observing individual plants for 1 minute and documenting the number of honey bees and non-*Apis* insects foraging. Observations were documented 3 times an hour throughout the day (900-1400) for one day. The average number of *Apis* and non-*Apis* visitors per plant and the total number of each visitor type observed was calculated for each site. For *Phacelia distans,* an additional site is included (DAW).

| **Plant spp.** | **Site** | **Total # of *Apis* Observed** | **Total # of non-*Apis* Observed** | **Avg # of *Apis* per plant** | **Avg # of**  **non-*Apis***  **per plant** | **% *Apis*** |
| --- | --- | --- | --- | --- | --- | --- |
| ***Salvia apiana*** | MTR | 386 | 16 | 7.72 | 0.32 | 0.96 |
| ***Salvia apiana*** | BSR | 230 | 19 | 4.6 | 0.38 | 0.92 |
| ***Salvia apiana*** | ECR | 203 | 21 | 4.06 | 0.42 | 0.91 |
| ***Salvia apiana*** | 56BP | 183 | 12 | 3.66 | 0.24 | 0.94 |
| ***Salvia mellifera*** | GC | 226 | 16 | 4.52 | 0.32 | 0.93 |
| ***Salvia mellifera*** | MTR | 334 | 6 | 6.68 | 0.12 | 0.98 |
| ***Phacelia distans*** | 56BP | 99 | 24 | 1.98 | 0.48 | 0.80 |
| ***Phacelia distans*** | DAW | 104 | 24 | 2.08 | 0.48 | 0.81 |

**Table S13. Study site locations.** ECR (Elliott Chaparral Reserve) and DAW (Dawson Los Monos Canyon Reserve) are University of California Natural Reserve System Reserves.

| **Plant spp.** | **Site** | **Latitude** | **Longitude** |
| --- | --- | --- | --- |
| ***Salvia apiana*** | MTR | 32.839967 | -117.067744 |
| ***Salvia apiana*** | BSR | 33.014405 | -117.012144 |
| ***Salvia apiana*** | ECR | 32.893315 | -117.088658 |
| ***Salvia apiana*** | 56BP | 32.935653 | -117.218702 |
| ***Salvia mellifera*** | GC | 32.965299 | -117.223841 |
| ***Salvia mellifera*** | MTR | 32.839967 | -117.067744 |
| ***Phacelia distans*** | 56BP | 32.935653 | -117.218702 |
| ***Phacelia distans*** | DAW | 33.151313 | -117.249102 |
